# Tryptophan wasting and disease activity as a systems phenomenon in inflammation – an analysis across 13 chronic inflammatory diseases

**DOI:** 10.1101/2023.08.25.554383

**Authors:** Danielle MM Harris, Silke Szymczak, Sven Schuchardt, Johannes Labrenz, Florian Tran, Lina Welz, Hanna Graßhoff, Henner Zirpel, Melike Sümbül, Mhmd Oumari, Nils Engelbogen, Ralf Junker, Claudio Conrad, Diamant Thaçi, Norbert Frey, Andre Franke, Stephan Weidinger, Philip Rosenstiel, Bimba Hoyer, Silvio Waschina, Stefan Schreiber, Konrad Aden

## Abstract

Chronic inflammatory diseases (CID) are systems disorders affecting various organs including the intestine, joint and skin. The essential amino acid tryptophan (Trp) is not only used for protein synthesis but can also be catabolized to various bioactive derivatives that are important for cellular energy metabolism and immune regulation. Increased Trp catabolism via the kynurenine pathway is seen across individual CID entities^1–5^. Here, we assessed the levels of Trp and tryptophan derivatives across 13 CID to investigate the extent and nature of Trp wasting as a systems phenomenon in CID. We found reduced serum Trp levels across the majority of CID and a prevailing negative relationship between Trp and systemic inflammatory marker C-reactive protein (CRP). Increases in the kynurenine-to-Trp ratio (Kyn:Trp) indicate that the kynurenine pathway is a major route for CID-related Trp wasting. However, the extent of Trp depletion and its relationship with disease activity varies by disease, indicating potential differences in Trp metabolism. In addition, we find that amino acid catabolism in chronic inflammation is specific to tryptophan wasting, whereas other proteinogenic amino acids are not affected. Hence, our results suggest that increased Trp catabolism is a common metabolic occurrence in CID that may directly affect systemic immunity.

**Grant support:** This work was supported by the DFG Cluster of Excellence 1261 “Precision medicine in chronic inflammation” (KA, SSchr, PR, BH, SWa), the BMBF (e:Med Juniorverbund “Try-IBD” 01ZX1915A and 01ZX2215, the e:Med Network iTREAT 01ZX2202A, and GUIDE-IBD 031L0188A), DFG RU5042 (PR, KA), and Innovative Medicines Initiative 2 Joint Undertakings (“Taxonomy, Treatments, Targets and Remission”, No. 831434, “ImmUniverse”, grant agreement No. 853995, “BIOMAP”, grant agreement No. 821511).

## Main

To investigate the role of Trp metabolism in CIDs, we implemented serum Trp measurements into clinical laboratory assessments at our CID outpatient clinic (UKSH, Kiel, Germany). This cohort (cohort 1, Fig. 1A) compromises data from patients with diseases affecting the gastrointestinal tract (GI), musculoskeletal (MSK) system, and skin. GI CIDs included Crohn’s disease (CD) and ulcerative colitis (UC). MSK CIDs included diseases affecting joints (seronegative RA: RAneg, seropositive RA: RApos, axial spondyloarthritis: AxSpA, psoriatic arthritis: PsA), connective tissue (SLE, systemic sclerosis: SSc, Sjögren’s syndrome: SjS, unspecified connective tissue disease: CTD), and blood vessels (giant cell arteritis: GCA, polymyalgia rheumatica: PMR). Finally, cohort 1 includes psoriasis (Pso), which primarily affects the skin. Cohort 1 includes longitudinal measurements of 1949 patients with a total number of 29,012 observations, and an average number of 14.7 ± 16.6 (mean ± standard deviation) follow-ups. Additionally, single observations of 291 healthy individuals are included as a reference (Supplementary Table 1). Serum Trp was reduced in 9 of the 13 CIDs tested, with no detectable reduction in serum Trp CTD, AxSpA, PsA or Pso patients (Fig. 1B). Considering the potential influence of disease activity on the patterns observed, we used the established systemic inflammation marker CRP as a proxy for disease activity. Using a cut-off value of 5 mg/L to designate inactive disease, we identified serum Trp reduction in a smaller selection of CIDs even during periods of biochemical disease inactivity: specifically, in CD, UC, RApos, SjS, and SLE (Fig. 1C). To assess the relationship between disease activity and Trp levels, we analysed the association between CRP and Trp using linear mixed models for each CID. We observed significant negative relationships between serum CRP and Trp in most of the CID studied, except for CTD and SjS, which showed non-significant negative trends (Fig. 1D). In summary, our findings suggest that decreased Trp levels are associated with systemic inflammatory activity across various CID, which is consistent with the previously described role of Trp in inflammatory processes^4^. Furthermore, excessive serum Trp reduction in the absence of overt systemic inflammation is a feature of a smaller subset of CID, indicating that Trp may indicate subtle immune activation in the absence of overt inflammation.

**Fig. 1.**
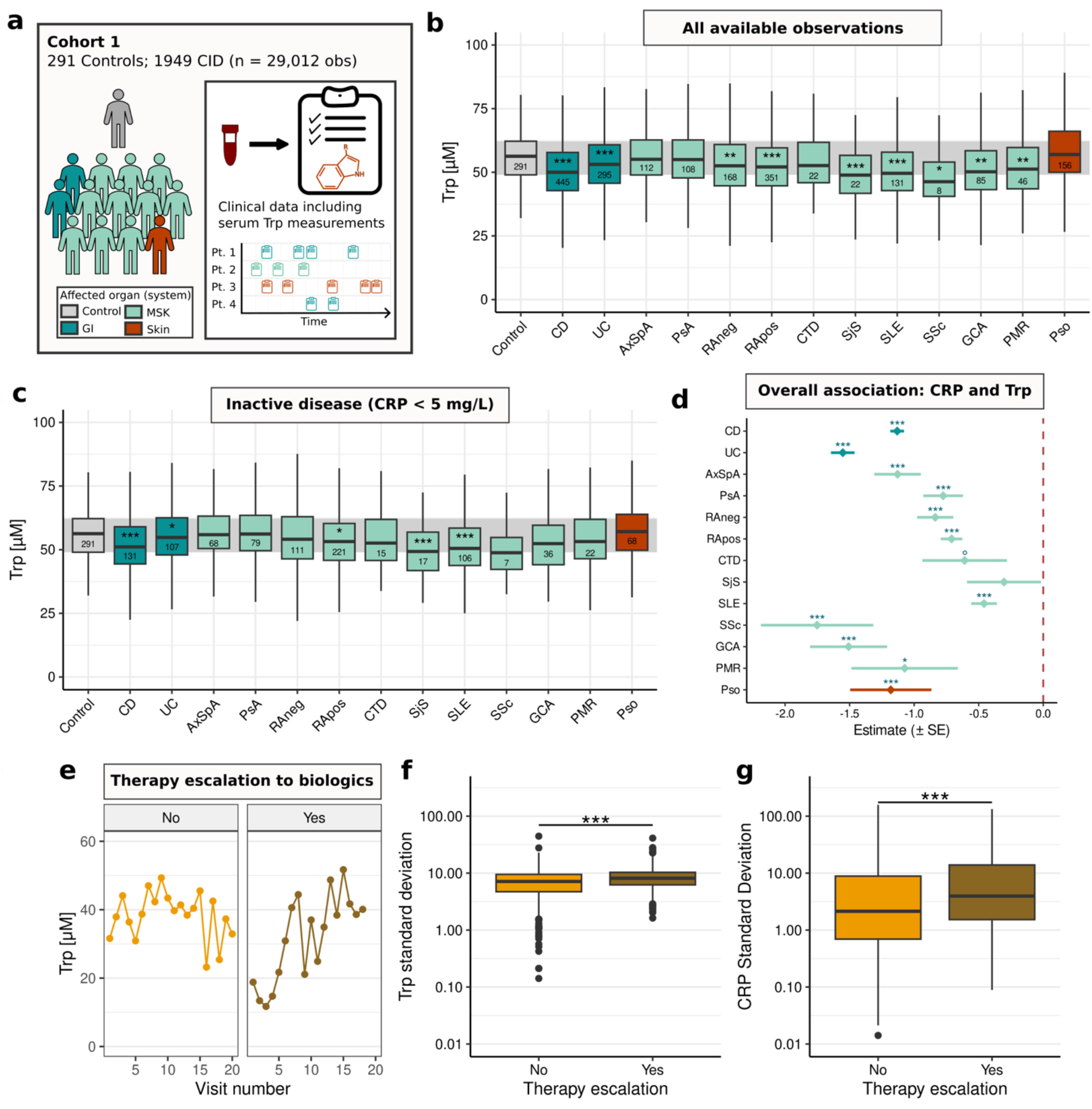
Reduced serum Trp across CIDs. **a**, Cohort consists of 13 different CIDs plus a healthy reference group. **b**, Assessment of serum Trp levels across all available observations. Here and in the next panel, numbers within the boxes indicate the number of patients included in the analysis and asterisks indicate significance in comparison with controls. **c**, Serum Trp levels of CID patients using only serum samples from patients with low disease activity, defined as CRP < 5 mg/L. **d**, Linear mixed models were used to assess the relationship between CRP and Trp for each CID. Estimates depict the estimated change to CRP associated with a one-unit change in Trp (both variables were log_10_ transformed). **e**, Exemplary Trp trajectories for two patients; one requiring biologic therapy escalation and one who did not. **f**,**g** Within-patient standard deviations in Trp and CRP levels are higher for patients undergoing therapy escalation compared to those who did not. Significance annotation: ° FDR < 0.1; * FDR < 0.05, ** FDR < 0.01, *** FDR < 0.001.

We further hypothesised that disrupted Trp metabolism in CID is indicative for the long-term disease course. Critical transition theory posits that increased variance is an indicator of a shift in previously stable systems^6^. This holds true for diabetes, where long-term glycaemic variability is associated with cardiovascular complications^7^. We adapted this concept for Trp metabolism to investigate whether Trp variability indicates complicated disease course in CID. Targeted immunomodulatory therapies (such as antibodies against cytokines or kinase inhibitors; hereafter referred to as biologics) are commonly used as second-line treatment upon failure of conventional medications (e.g., disease-modifying antirheumatic drugs, topical agents, corticosteroids, aminosalicyclates). Therefore, patients not requiring biologics during the observation period are expected to experience less aggressive disease courses compared to patients who undergo therapy escalation. Therefore, the fact that patients requiring biologic therapy escalation had higher variation in Trp and CRP levels compared to patients not requiring escalation (Fig. E-G), is evidence that fluctuating serum Trp levels are linked to disease course severity. Thus, in line with critical transition theory, CRP and Trp variability may indicate phase shifts toward more complicated diseases courses, suggesting possible utility for these metrics in a predictive context.

Next, we assessed links between serum Trp levels and available clinical disease activity indices. Consistent with previous reports^8^, we confirmed a negative association between Crohn’s disease activity index (CDAI) and total Mayo scores in CD and UC (n = 87 and 96, respectively, Fig. 2AB). We further analysed the association between serum Trp and the Bath Ankylosing Spondylitis Disease Activity Index (BASDAI, n = 57 AxSpA patients), Disease Activity Score 28-CRP (DAS28, n = 59 RAneg and 134 RApos patients) and Psoriasis Area and Severity Index (PASI, n = 22). We identified significant negative associations between BASDAI and Trp (Fig. 2C), as well as between DAS28 and Trp for RAneg (Fig. 2D) but, most surprisingly, not for RApos (Fig. 2E). There was no clear relationship between PASI and Trp (Fig. 2F). These differences in the strength of the association between Trp and disease activity indices could potentially indicate systemic differences in immune network perturbation. It is also possible that the total inflammatory burden was not high enough to reveal an association between PASI, which solely evaluates skin symptoms, and Trp.

**Fig. 2.**
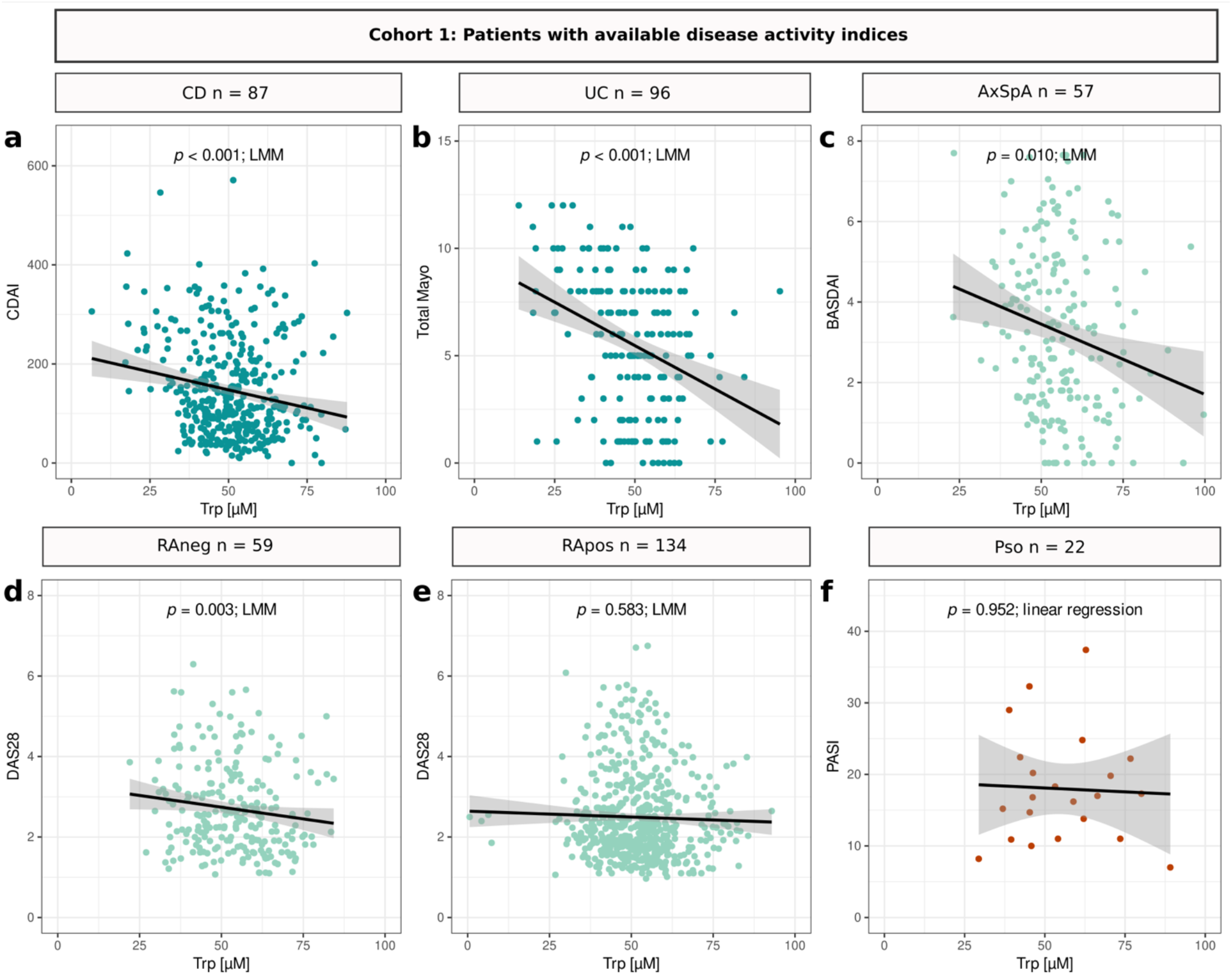
Association between Trp and different disease activity indices in cohort 1. **a-c**, Negative relationships between Trp and CDAI, total Mayo, and BASDAI for CD, UC, and AxSpA, respectively. **de**, The relationship between DAS28 and Trp in RA varies by disease entity, with a detectable negative association between DAS28 and Trp in RAneg but not RApos patients. **f**, No significant association between Trp and PASI was detected in Pso patients. All available observations are included in each plot, and the regression line is depicted in black. LMM: Lineaer mixed model, n = total number of patients.

To identify the primary driver of Trp reduction, we considered four potential contributors to serum Trp levels: (1) dietary protein uptake, (2) protein synthesis, (3) targeted Trp degradation (e.g., via the kynurenine pathway), and (4) microbial influences via dietary Trp degradation or de novo Trp production. Accordingly, we selected patients with extreme Trp levels for inclusion into cohort 2; these patients had Trp values below 45 μM or above 60 μM. We analysed Trp and selected Trp derivatives plus canonical proteogenic amino acids via targeted metabolomics with serum samples from CID patients (58 GI, 88 MSK and 52 Skin) and 179 healthy individuals (Fig. 3AB, Supplementary Table 2). In addition, we assessed faecal Trp levels through targeted metabolomics from a sub-cohort of 164 patients (51 GI, 67 MSK and 46 Skin). We found no evidence of differing faecal Trp levels in CID patients with high versus low serum Trp levels, neither at the level of the affected organ group, nor when the diseases were combined (Fig. 3C), suggesting no longstanding microbial influence on Trp levels. However, it must be acknowledged that these stool samples were not obtained on the date of serum sampling (mean difference -31.3 days) so we cannot rule out short term changes to faecal Trp levels. Considering other potential drivers of Trp depletion, in cases of dietary protein restriction, reduced uptake or increased protein synthesis, Trp should be reduced along with all or specific proteogenic amino acids. Conversely, under excessive Trp catabolism, we expect Trp levels to be largely independent from other amino acid levels since they are not required for Trp-specific catabolic processes. Thus, we correlated Trp with 18 other proteogenic amino acids within three groups: healthy individuals, and CID patients with low and high serum Trp.

**Fig. 3.**
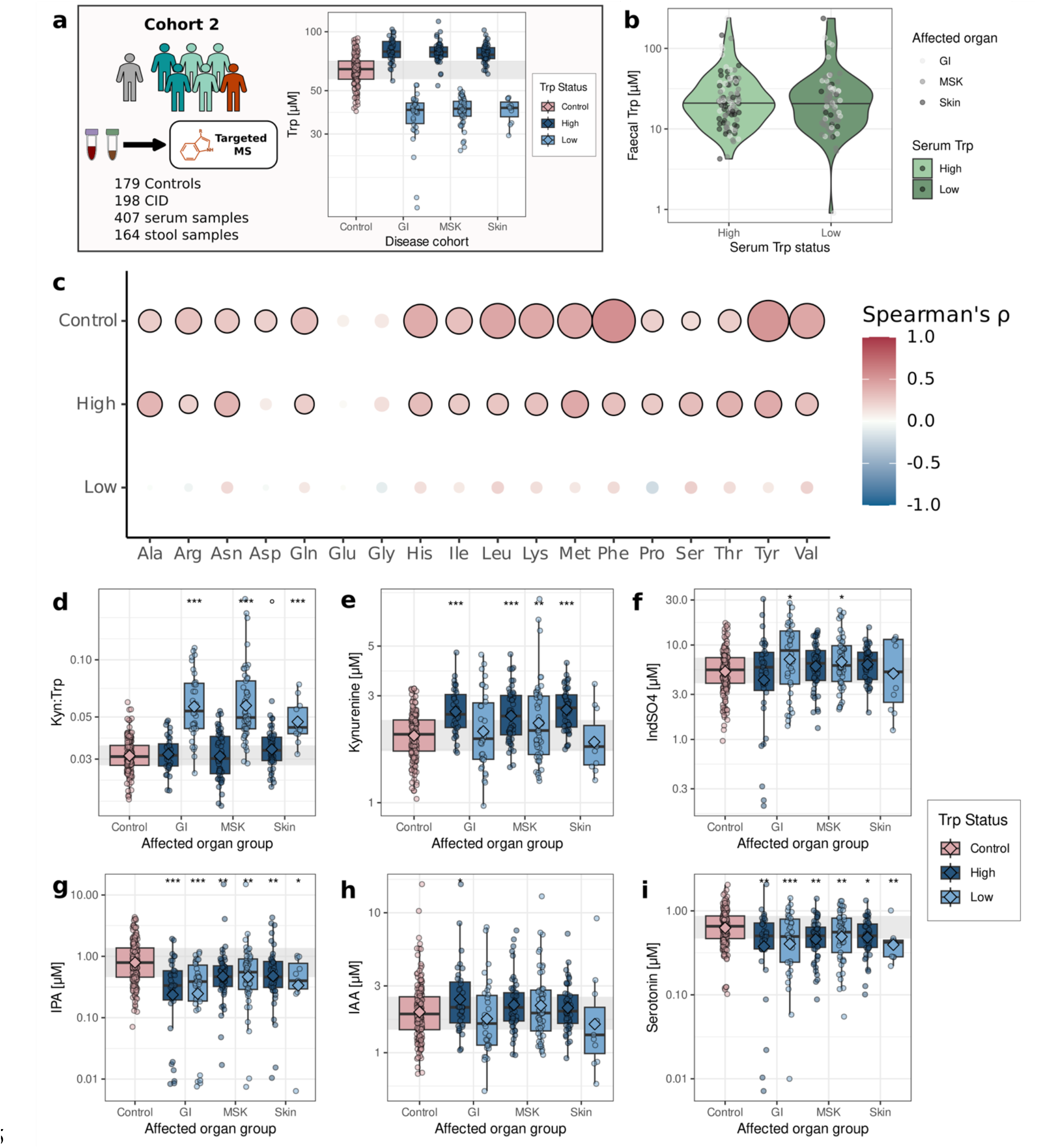
Lower serum Trp levels accompany kynurenine-pathway activation. **a**, Cohort 2 overview: serum samples from 198 CID patients plus 179 controls were selected for mass spectrometry-based targeted metabolomics. Additionally, 164 stool samples from CID patients were sent for metabolomics analysis. Patients were selected for cohort 2 based on available HPLC-based Trp measurements (below 45 μM and above 60 μM). Mass spectrometry-based serum Trp measurements are included here as an overview. **b**, Faecal Trp levels in high and low serum Trp groups. **c**, Partial Spearman’s correlations between Trp and the other measured proteogenic amino acids. Correlations with FDR < 0.05 are circled in black, with increasing circle size denoting decreasing FDR via calculation of -log10(FDR). **d-i**, Serum levels of selected Trp derivatives and the ratio of kynurenine-to-Trp (Kyn:Trp). Diamonds indicate means and comparisons are made against the control group. Significance annotation: * FDR < 0.05, ** FDR < 0.01, *** FDR < 0.001.

As anticipated, we observed strong, positive correlations between Trp and 16 of 18 amino acids in healthy individuals, suggesting that under physiological conditions, serum Trp concentration fluctuates with dietary levels, general amino acid uptake and/or protein synthesis (Fig. 2C). CID patients with high serum Trp had notably intact correlation networks compared to CID patients with low serum Trp (15 versus 0 amino acid correlations with Trp, respectively) (Fig. 2C). Increased correlations in CID patients with high Trp compared to patients with low Trp may imply that elevated Trp levels resulted from increased amino acid intake or decreased protein synthesis. Moreover, zero amino acids correlated with Trp in CID patients with low serum Trp. This implies an increased influence of Trp catabolism on serum Trp levels in low Trp CID patients relative to the influence of diet or protein synthesis. It also underlines a unique role of the catabolism of tryptophan, among other amino acids, in the context of systemic inflammation. Consistent with this theory, CID patients with high Trp values had increased levels of 13 amino acids relative to the healthy population, suggesting that in CID, extremely high serum Trp arises due to differences in protein intake or use. Conversely, CID patients with low serum Trp have more variation in amino acid profiles, with half of amino acids either higher or not significantly compared to healthy individuals (n = 2 and 7, respectively, Supplementary Fig. 1). Extending the correlation networks to all measured proteogenic amino acids reflects this volatility, with notably intact correlation networks for healthy individuals and CID patients with high Trp, and less overall correlations between amino acids for CID patients with low Trp (Supplementary Fig.2).

Next, we quantified representative metabolites from the three main Trp catabolic routes: the kynurenine pathway, serotonin pathway and microbial indole production^1^. We focus our discussion on Trp derivatives that are higher in low Trp CID patients relative to healthy individuals since they are most likely to convey information about the fate of Trp under extreme catabolism rather than diet-specific information. The implicit assumption here is that metabolites accumulate when Trp is catabolized along the respective pathways, providing information about which pathways may actively degrade Trp.

Trp Degradation via the kynurenine pathway results in the production of a variety of immunomodulatory metabolites^2^. The rate-limiting step of this pathway is the production of kynurenine itself, thus, Kyn:Trp is often used as a proxy for the pathway’s activity^9^. Our analysis revealed that Kyn:Trp is highest in low Trp CID patients compared to healthy individuals across all CID groups (Fig. 3D). We further assessed the levels of kynurenine alone, which were elevated in all disease entities when Trp was high and, strikingly, even when Trp levels were low in MSK CID (Fig. 3E). Hence, our data strongly indicate that reduced Trp in CID arises due to Trp wasting along the kynurenine axis in all investigated CID.

Out of the four other Trp derivatives measured (indoxylsulfate (IndSO4), indole-3-acetic acid (IAA), indole propionic acid (IPA), and serotonin), IndSO4 was the only derivative with increased levels in low Trp CID patients relative to healthy individuals (Fig. 3F-I). On the contrary, relative to healthy individuals, IPA and serotonin were reduced in across CID in patients with low and high serum Trp (Fig. 3GI). This might point towards a general decrease in Trp metabolism along these catabolic routes, potentially because of increased kynurenine pathway demand.

CID share several pathophysiological commonalities beyond mere tissue inflammation and destruction at sites of anatomical manifestation, e.g., chronic kidney damage or fatigue^10,11^. In this comprehensive analysis, we tested the hypotheses that Trp reduction is common in CID and that this reduction arises due to increased kynurenine pathway activity. Doing so, we identified a consistent negative association between serum Trp and CRP levels. In the light of our targeted metabolomics analysis, these findings indicate that Trp wasting along the kynurenine axis is an overarching phenomenon of acute flares of chronic inflammation. In this context, increased metabolic demand along the kynurenine pathway upon high systemic inflammatory load is likely a major contributor to decreasing serum Trp levels. Increased Trp degradation via this pathway has been observed in numerous cell types as a result of inflammatory cytokine signalling (e.g., interferon gamma and transforming growth factor beta)^12,13^. In several cancer types, upregulation of the kynurenine pathway is generally considered as a hallmark of immune evasion as it may suppress anti-tumour immunity^12^. This suggests that Trp catabolism could be upregulated in CIDs due to aberrant cytokine signalling without direct proinflammatory consequences or even as a futile attempt to down-regulate excessive immune signalling. Therefore, consideration of possible pathological connections between excessive kynurenine pathway activity in CID needs to be weighed against the fact that this pathway is also active in immunosuppressive environments (i.e., cancer). Harden et al. proposed a model that would explain this apparent dichotomy^6^. The model contrasts the enzymes involved in the first step of the kynurenine pathway (indoleamine 2,3-dioxigenase 1 (IDO1) and tryptophan dioxygenase (TDO)) with downstream kynureninase expression. Kynureninase outweighs combined expression of IDO and TDO in CID, while the opposite pattern occurs in cancer. Hence, CID should have an enrichment of downstream proinflammatory metabolites with a relative paucity of upstream anti-inflammatory metabolites. Although we have extended our findings to include more CID, kynureninase predominance over IDO-TDO activity could explain the generalizability of low Trp across CID observed here and is suggestive that under the right conditions, excessive kynurenine pathway activity may contribute to proinflammatory environments.

Most importantly we were able to identify a subgroup of disease entities (CD, UC, RApos, SjS, SLE) with reduced Trp levels even in absence of systemic inflammation (CRP < 5 mg/L). As our data indicate that Trp reduction results from enzymatic degradation, these disease entities may have distinct inflammatory metabotypes in which subtle Trp degradation is an initial causal pathophysiological event. Interestingly, the induction of indoleamine 2,3-dioxygenase (IDO), the enzyme responsible for Trp degradation into kynurenine^2^, is expressed in a Janus kinase-signal transducer and activator of transcription (JAK-STAT) dependent manner^14^. Overactivation of the JAK-STAT pathway is present in some CID^15–17^ and inhibition of this pathway is therapeutically employed using pan-JAK or JAK1-selective inhibitors in CD^18^, UC^19^, RA^15,20^, and SLE^17^. Hence, our data suggest that monitoring of Trp metabolism in disease entities where Trp levels tend to be low might not only be a sensitive and precise biomarker but could facilitate a more targeted therapeutic approach in CID.

We also found increased IndSO4 in GI and MSK CID patients with low Trp. Inflammatory effects of IndSO4 have been investigated in chronic kidney disease, where it disrupts the intestinal barrier and contributes to inflammation via activation of the nuclear factor kappa B signalling pathway^22^. Of note, decreased kidney function is a potential complication of GI and MSK CID^23,24^. Two other Trp derivatives displayed notable trends relative to the healthy population tested: serotonin and IPA. Studies on serum serotonin and CID have reported conflicting results, e.g., in one study, RA patients had lower serum serotonin relative to the healthy population^21^, while another reported higher levels^25^. Administration of anti-tumour necrosis factor drugs do reportedly lower serum serotonin levels^26^, which may be contributing to the reported differences ^21,27,28^. IPA has anti-inflammatory effects that are mediated via aryl hydrocarbon receptor (AHR) signalling and its serum levels are reportedly reduced in UC in a disease activity-dependent manner^29,30^. We identified IPA reduction across CID, indicating a paucity of AHR signalling as a potential joint feature of CID.

Important limitations of our study include the fact that serum Trp measurement was conducted in a cross-sectional cohort, so we cannot draw conclusions about metabolic dynamics that arise before or after successful treatment in individual patients. Although our targeted metabolomics data strongly indicate active enzymatic degradation of Trp along the kynurenine pathway, the co-occurrence of metabolites along one metabolic pathway (Kyn:Trp) does not specifically measure the degree of Trp wasting. Hence, dynamic measurement of Trp degradation (e.g., in-vivo Trp fluxomics^31^) should be considered as a novel diagnostic element in CID patients. This would not only help to discern the individual degree of Trp wasting but also help uncover the metabolic fate of Trp. The approach would additionally help overcome the lack of detailed dietary information, which also critically impacts host Trp levels and cannot be completely disentangled within cross-sectional cohorts. Taken together, our results indicate that excessive kynurenine pathway activity occurs across CID, but the extent and circumstances leading to activation of the pathway may have disease-specific aspects. We therefore suggest that a detailed exploration of Trp turnover in CID patients might provide a deeper insight into the molecular origins of disease pathophysiology and might also lead to different therapeutic concepts for targeted therapies.

## Methods

### Patient recruitment

#### Cohort 1

In the IBD outpatient clinic of the Department of Internal Medicine I at the University Medical Center Schleswig-Holstein (Campus Kiel, Germany) and adult patients with confirmed diagnosis of CD, UC, RAneg, RApos, SLE, SSc, PMR, AxSpA, GCA, CTD, Pso, and PsA were included in retrospective analysis of clinical phenotyping and clinical laboratory diagnostics (Supplementary Table 1). In total 1949 patients were recruited, with routine laboratory assessment at every clinical visit, including serum Trp measurements via high performance liquid chromatography (HPLC) measurement (detailed below). Clinical visits spanned from January 2015 to June 2023. In addition, 291 healthy controls were recruited, resulting in a total of 29,012 clinical observations.

#### Cohort 2

Patient recruitment occurred at the IBD outpatient clinic of the Department of Internal Medicine I at the University Medical Center Schleswig-Holstein Kiel Campus, and the Comprehensive Center for Inflammation Medicine and Clinic of Rheumatology at the University Medical Center Schleswig-Holstein Lübeck Campus in Germany. This multi-CID cohort included patients recruited in 2020 with UC, CD, Pso, RApos, RAneg, among others (Supplementary Table 2). Patients were included based on HPLC-based Trp measurement. Selection was based on serum Trp values below 45 μM or above 60 μM (as measured by HPLC). In total, the cohort consists of 407 serum samples from 198 patients. In addition, serum metabolomes from 179 healthy individuals were analysed. This cohort was recruited at the University Hospital Schleswig Holstein, Campus Kiel 2016 and comprised detailed phenotypic, disease related and dietary information. The study was approved by the local ethic committee in Kiel (D441). None of the participants had received any antibiotics or other medication 2 months prior to inclusion and individuals with diabetes were excluded from the analysis.

#### Ethics

Approval was granted by the ethics committee of the medical faculty of Kiel University before the start of the study (AZ D489/14).

### Measurement of serum metabolites

#### HPLC

Serum levels of Trp were determined using an In Vitro Diagnostic Conformité Européenne (CE) certified HPLC kit. Lower limits of quantification and lower limits of detection were calculated according to DIN 32645 guidelines.

#### Targeted Mass Spectrometry of Trp derivatives

Trp derivatives were measured in the patient serum and stool of cohort 2 via tandem mass spectrometry (LC-MS-MS) using the MxP Quant 500 kit (Biocrates Life Sciences AG, Innsbruck, Austria) according to the manufacturer’s instructions. Serum samples were collected using serum s-monovette (9 mL, Sarstedt, Germany) and incubated upright at room temperature for 30 minutes prior to a 10-minute centrifugation at 2000*g*. Serum was aliquoted in 500 μL tubes and stored at -80°C until sample preparation for MS. The MxP Quant 500 kit simultaneously measures 630 metabolites covering 14 small molecule and 12 different lipid classes. It combines flow injection analysis tandem mass spectrometry (FIA-MS/MS) using SCIEX 5500 Q-Trap mass spectrometer (SCIEX, Darmstadt, Germany) for lipids and liquid chromatography tandem mass spectrometry (LC-MS/MS) using Agilent 1290 Infinity II liquid chromatography (Santa Clara, CA, USA) coupled with a SCIEX 5500 Q-Trap mass spectrometer for small molecules using multiple reaction monitoring (MRM) to detect the analytes. Data evaluation for serum metabolite concentrations and quality assessment was performed with the software SCIEX Analyst software (Version 1.7.2) and the MetIDQ™ software package (Oxygen-DB110-3023), which is an integral part of the MxP Quant 500 kit.

## Data analysis

### Pre-processing of metabolomics data

Metabolite concentrations below the lower limit of quantitation were removed from the analysis (i.e., considered missing values), and metabolites were further filtered based on the 80% rule^32^: only metabolites with 80% non-missing values were included for further analysis. The remaining missing values were imputed using GSimp^33^, which uses iterative Gibbs sampling for missing value imputation. Only Trp, Trp derivatives, and proteogenic amino acids were considered for further analysis.

### Statistical analysis

To assess the difference between the serum Trp (derivative) concentrations in controls and different CID entities as well as in patients with high and low serum Trp, we used linear mixed models with Trp as the dependent variable, and CID entity and sex as fixed effects, with patient IDs as random intercepts. Because biological sex influences Trp levels^34^, it was included as a fixed effect in all models. Assessment of differences in serum amino acids between control and high/low Trp groups included the primary affected organ or organ system (i.e., GI, MSK and Skin) as an additional fixed effect, i.e., metabolite ∼ Trp status + disease cohort + sex + (1 | Patient). The reference group in the aforementioned cases were the healthy individuals. To assess the predictive power of Trp on CRP and disease activity indices, Trp was modelled as a fixed effect (predictor variable) with the disease activity metric as the dependent variables (e.g., CRP ∼ Trp + sex + (1 | Patient)). Models were fit using maximum likelihood (REML) and the degrees of freedom were estimated using Satterwaithe’s method. Linear models were employed to estimate the relationship between Trp and PASI, without an adjustment for biological sex as there was only one female patient with available Trp and PASI. Each linear (mixed) model was checked for violations in linear regression assumptions via visual inspection of fitted versus residual plots. The metabolites, CRP, as well as DAS28 scores were all log_10_ transformed. Partial correlations included an adjustment for biological sex and in order to meet the assumption of sample independence, only the first observation per patient was used in this analysis. A Student’s t-test was used to assess differences between faecal amino acid levels in the high and low serum Trp groups. Previous reports have demonstrated low Trp levels in some CIDs, so we specifically assessed whether the Trp levels were lower in the disease groups by calculating left-tailed p-values; all other tests were performed two-sided. We adjusted for multiple comparisons using the Benjamini-Hochberg procedure. In cohort 1, the false discovery rate correction was made across the different disease entities. In cohort 2, it was made across the metabolites and affected area for Trp high and Trp low groups together (i.e., across 36 p-values). To compare variation in Trp values between patients undergoing therapy escalation, patients the standard deviation of Trp and CRP was calculated at the patient level for patients with 3 or greater observations. Differences between groups were assessed using the Wilcoxon Rank Sum test. Analyses were performed with the statistical software R (version 4.2.1)^35^. R packages lme4 (version 3.1-3)^36^ and lmerTest (version 1.1-30)^37^ were used to fit the linear mixed models, and ppcor for partial Spearman correlations (version 1.1)^38^.

