## Supplemental Tables and Figures for "Tryptophan wasting and disease activity as a systems phenomenon in inflammation – an analysis across 13 chronic inflammatory diseases"

**Supplementary Table 1 | Characteristics of cohort 1.** Mean Trp level is reported across all available observations. UC: ulcerative colitis, CD: Crohn's disease, RApos: seropositive rheumatoid arthritis, RAneg: seronegative rheumatoid arthritis, SSc: systemic sclerosis, SLE: Systemic Lupus Erythematosus, PMR: polymyalgia rheumatica, GCA: giant cell arteritis, CTD: unspecified connective tissue disease, AxSpA: axial spondyloarthritis, Pso: Psoriasis, PsA: Psoriatic arthritis, SjS: Sjögren's syndrome. Mean Trp was calculated across all available observations.

| | **Control** | **UC** | **CD** | **RApos** | **RAneg** |
|---|---|---|---|---|---|
| No. individuals | 291 | 182 | 353 | 351 | 168 |
| No. observations | 291 | 3578 | 8223 | 5317 | 2567 |
| Trp (mean ± SD) | 56.58 (10.84) | 53.29 (13.05) | 50.32 (13.15) | 52.81 (12.35) | 53.49 (12.54) |
| Sex = Female/Male | 109/182 | 135/159 | 250/195 | 246/105 | 114/54 |
| Age (mean ± SD) | 38.16 (14.07) | 42.46 (15.37) | 44.52 (14.80) | 62.35 (12.58) | 63.90 (13.52) |
| Therapy escalation | | | | | |
| No | NA | 80 | 113 | 55 | 29 |
| Yes | NA | 82 | 214 | 143 | 69 |
| | **SSc** | **SLE** | **PMR** | **GCA** | **CTD** |
| No. individuals | 8 | 131 | 46 | 85 | 22 |
| No. observations | 232 | 3365 | 173 | 400 | 275 |
| Trp (mean ± SD) | 47.41 (10.64) | 50.85 (11.55) | 51.68 (11.93) | 52.22 (12.38) | 53.71 (10.44) |
| Sex = Female/Male | 05/03 | 124/7 | 26/20 | 59/26 | 19/3 |
| Age (mean ± SD) | 60.12 (11.69) | 53.41 (12.70) | 70.52 (8.43) | 69.45 (8.99) | 49.36 (12.14) |
| Therapy escalation | | | | | |
| No | 7 | 99 | 0 | 0 | 18 |
| Yes | 1 | 30 | 0 | 0 | 4 |
| | **AxSpA** | **Pso** | **PsA** | **SjS** | |
| No. individuals | 112 | 156 | 108 | 22 | |
| No. observations | 1287 | 845 | 2066 | 393 | |
| Trp (mean ± SD) | 56.24 (11.97) | 58.00 (12.21) | 55.96 (12.38) | 48.99 (9.61) | |
| Sex = Female/Male | 36/76 | 47/109 | 57/51 | 18/4 | |
| Age (mean ± SD) | 45.96 (13.04) | 51.29 (14.19) | 57.07 (12.45) | 62.27 (13.59) | |
| Therapy escalation | | | | | |
| No | 9 | 123 | 18 | 19 | |
| Yes | 35 | 10 | 73 | 3 | |

**Supplementary Table 2 | Characteristics of cohort 2.** Total patients in the gastrointestinal (GI) disease category were UC (16) and CD (40), indeterminate colitis (2). Patients with diseases affecting the musculoskeletal system (MSK) included RAneg (16), RApos (33), SSc (1), SLE (13), CTD (3), unspecified spondyloarthritis (15), unspecified rheumatoid arthritis (2), SjS (4), and polymyositis (1). Skin inflammatory disease was represented by Pso only.

| | **Control** | **GI** | **MSK** | **Skin** |
|---|---|---|---|---|
| No. individuals | 179 | 58 | 88 | 52 |
| No. serum samples | 179 | 69 | 101 | 58 |
| No. stool samples | NA | 51 | 67 | 46 |
| Trp (mean ± SD) | 64.71 (11.14) | 59.80 (23.74) | 60.24 (21.31) | 70.43 (16.87) |
| Sex = Female/Male | 80/99 | 33/25 | 57/31 | 22/30 |
| Age (mean ± SD) | 46.58 (8.20) | 49.74 (15.82) | 60.47 (12.92) | 54.85 (12.77) |
| Trp status (%) | | | | |
| Control | 179 (100.0) | NA | NA | NA |
| High | NA | 28 (48.3) | 45 (51.1) | 43 (82.7) |
| Low | NA | 30 (51.7) | 43 (48.9) | 9 (17.3) |

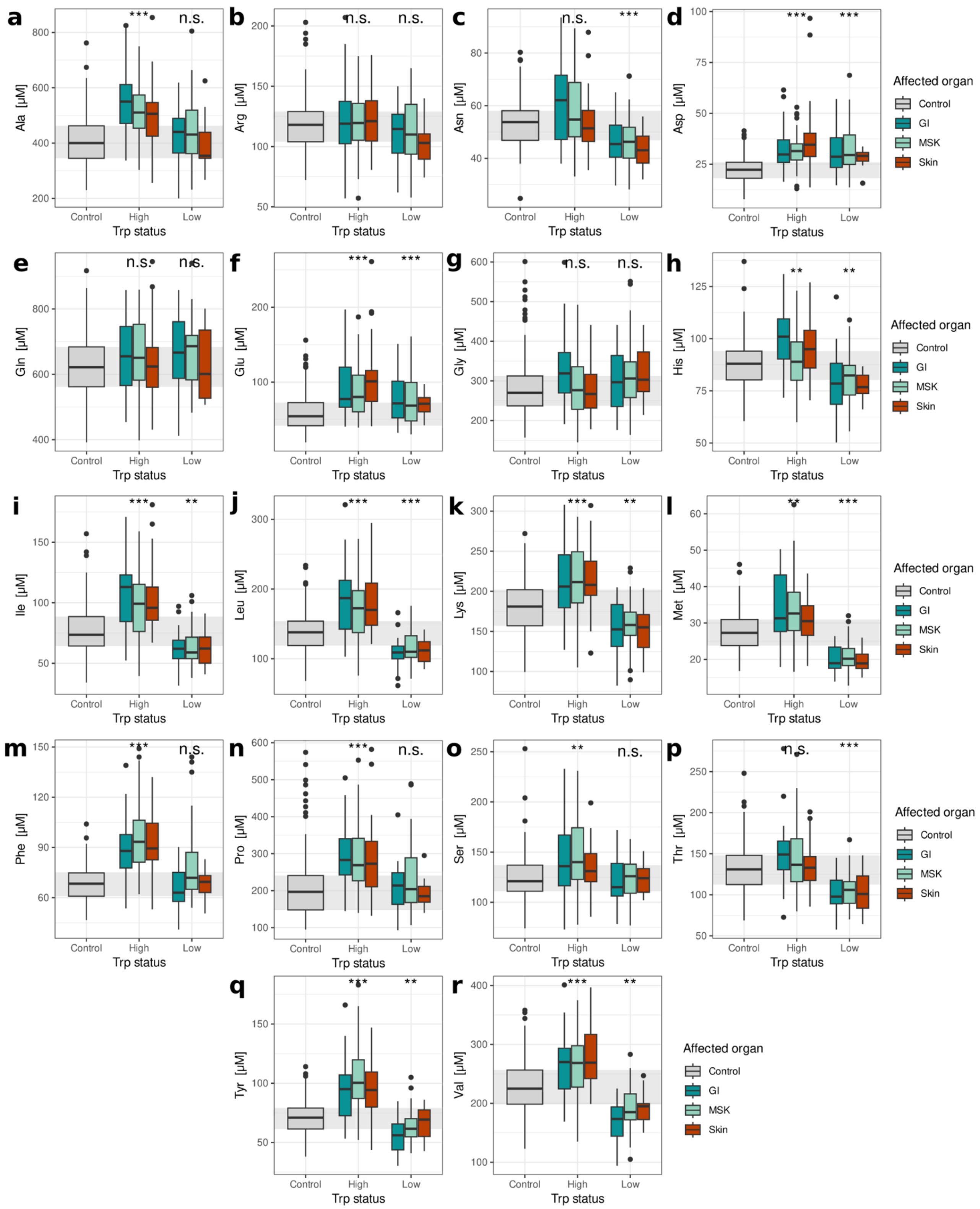


**Supplementary Figure 1 | Levels of serum amino acids in the serum of patients with high and low serum Trp.** Ala: Alanine, Arg: Arginine, Asn: Asparagine, Asp: Aspartic acid, Gln: Glutamine, Glu: Glutamate, Gly: Glycine, His: Histidine, Ile: Isoleucine, Leu: Leucine, Lys: Lysine, Met: Methionine, Phe: Phenylalanine, Pro: Proline, Ser: Serine, Thr: Threonine, Tyr: Tyrosine, Val: Valine.

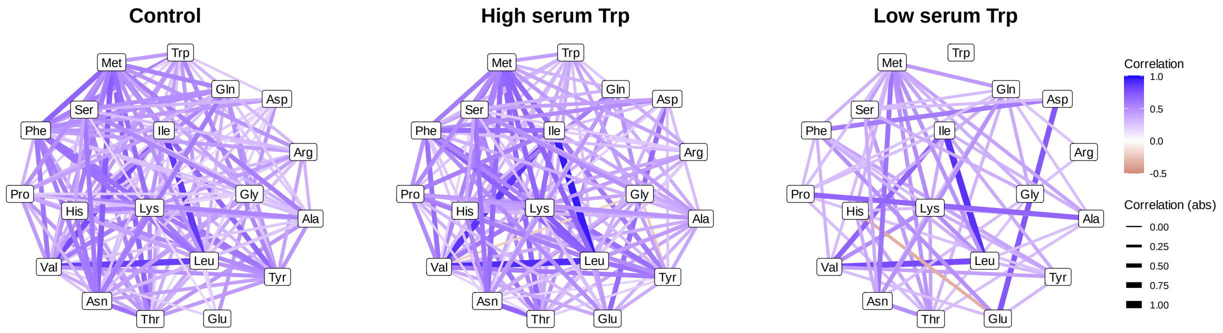


**Supplementary Figure 2 | Partial correlation networks between measured proteogenic amino acids in control individuals and CID patients with low and high serum Trp values.** Only correlations with FDR > 0.05 are plotted. Ala: Alanine, Arg: Arginine, Asn: Asparagine, Asp: Aspartic acid, Gln: Glutamine, Glu: Glutamate, Gly: Glycine, His: Histidine, Ile: Isoleucine, Leu: Leucine, Lys: Lysine, Met: Methionine, Phe: Phenylalanine, Pro: Proline, Ser: Serine, Thr: Threonine, Tyr: Tyrosine, Val: Valine.
